# WebCalEM: a browser-based tool for routine and accurate pixel size calibration in cryo-EM

**DOI:** 10.64898/2026.05.26.726020

**Authors:** Lan Dang, Zhiqing Wang, Sung Hyun (Joseph) Cho, Siduo Li, Geetanjali Chakraborty, Nahian Fyrose Fahim, Wen Jiang

## Abstract

Accurate determination of the image pixel size is critical for quantitative cryo-electron microscopy analyses, yet existing calibration methods remain under-utilized because installation barriers and workflow complexity discourage routine adoption. To fill in this gap, a web-based application, WebCalEM, was developed to transform specialized calibration procedures into an accessible routine practice. Micrographs of any specimen with a known crystalline lattice, such as gold or graphene-oxide, are uploaded through a standard browser, processed entirely client-side, and analyzed with real-time visualization and downloadable statistical outputs. The application is delivered as a single self-contained HTML file that runs in any modern web browser without server-side computation, a configuration well suited to isolated core-facility microscope workstations. Cross-standard consistency between gold and graphene-oxide measurements across two microscopes and ten magnification settings yields a Bland–Altman bias of −0.005% of nominal with 95% limits of agreement of [−0.30%, +0.29%]. By delivering this workflow with no local installation, WebCalEM lowers the practical barrier to documented per-dataset magnification calibration in routine cryo-EM operation.

**Synopsis:** WebCalEM is a browser-based, install-free application that performs routine cryo-EM pixel-size calibration directly from gold or graphene-oxide reflections in standard sample-support grids using sub-pixel Fourier-space peak localization; it reproduces the precision of established command-line calibration tools while removing the installation barrier and supporting retrospective per-region calibration on archived datasets.

## 1. Introduction

Magnification calibration to accurately determine the pixel size of a cryo-electron microscopy (cryo-EM) image dataset is critical for downstream biological structure determination (Tiwari *et al*., 2020; Dickerson *et al*., 2024; Wasilewski *et al*., 2012). At the high resolutions now standard for modern cryo-EM, errors of even ∼1% in pixel size compromise CTF phase fitting (Rohou & Grigorieff, 2015; Heimowitz *et al*., 2020), leading to incorrect spatial frequency determination and loss of information at high frequencies, ultimately limiting achievable resolution of the final structure (Dickerson *et al*., 2024). This issue is particularly important for amyloid reconstructions, where pixel-size errors cause the inferred cross-β spacing and helical rise to deviate from the expected 4.75 Å periodicity(Lövestam & Scheres, 2022; Scheres, 2020). More importantly, accurate calibration directly affects atomic modelling: even small scaling errors translate to incorrect atomic positions in the final structure(Tiwari *et al*., 2020; Dickerson *et al*., 2024; Wang *et al*., 2022). Accurate calibration is also required to place structures on a common absolute scale across instruments, time points, and facilities, and to report resolution in Å without over- or under-stating the true information content of a map (Penczek, 2010, 2020).

Three classes of methods are currently used for pixel-size calibration. External standards such as diffraction-grating replicas and Mag*I*Cal grids cover only narrow magnification ranges and can be biased by handling-induced distortion (McCaffrey & Baribeau, 1995; MAG*I*CAL™ Calibration Standard for TEM). Post-hoc calibration by atomic-model fitting against reference proteins such as apoferritin requires a full reconstruction pipeline and yields a single value per dataset, precluding real-time use or per-region statistics(Tiwari *et al*., 2020; Wasilewski *et al*., 2012; Wang *et al*., 2022). Internal crystalline references—gold {111} at 2.35 Å on UltrAuFoil/HexAuFoil supports (Russo & Passmore, 2014; Naydenova *et al*., 2020; Naydenova & Russo, 2022) and graphene/graphene-oxide reflections at 2.13 Å on standard supports (Han *et al*., 2020; Palovcak *et al*., 2018; Meyer *et al*., 2007; Sader *et al*., 2013)—provide directly measurable lattice spacings on every micrograph and span the magnification range relevant to single-particle and tomography cryo-EM.

While crystalline-reference calibration is the most promising route, existing implementations(Dickerson *et al*., 2024) are still largely standalone or offline tools rather than integrated components of routine cryo-EM workflows. Users typically install software locally, transfer data between packages, compile results from multiple measurements, and perform statistical analyses on the result separately. This fragmented workflow increases complexity, creating a substantial gap between available precise methods and their practical adoption. To close this practical gap, we present WebCalEM, a browser-based application that supports routine, SOP-compliant pixel size calibration. Users upload micrographs, interactively select regions of interest, and receive sub-pixel-precision pixel sizes from gold or graphene-oxide reflections. Optional ellipse fitting is exposed as an interactive diagnostic for users investigating projector-lens anisotropy or specimen tilt. WebCalEM supports manual and automated workflows, aggregates results across measurements with mean and standard deviation, and runs entirely in a standard web browser through client-side computation, lowering the practical barrier to routine, documented calibration on any computer and operating system.

## 2. Materials and methods

### 2.1. Grid preparation and EM data acquisition

UltrAuFoil gold grids were used as test samples for evaluating crystalline internal calibration standards. To introduce an additional crystalline reference, grids were coated with graphene oxide (GO) using a Langmuir–Blodgett-style deposition method (Palovcak *et al*., 2018). Data were collected on two microscopes at The Huck Cryo-EM Facility (RRID: SCR_0244456) at Penn State University: Thermo Fisher 200 kV Talos Arctica G2 and 300 kV Titan Krios G3. Micrographs were acquired at five nominal magnifications on each instrument (Arctica: 120,000×–310,000×, 0.36–0.97 Å/pixel; Krios: 105,000×–270,000×, 0.37–0.96 Å/pixel, defocus ranges: −0.5 to −2.0 µm). Exposure doses were 50–65 e^−^/Å^2^.

### 2.2. Image analysis and pixel size calculation

From 4K × 4K micrographs, regions of 1K × 1K to 2K × 2K pixels were transformed to reciprocal space by 2D fast Fourier transform (FFT). Calibration materials produce reflections at known spatial frequencies: graphene/ graphene oxide (GO) at 1/*d*_ref_ = 0.469 Å^-1^ (*d*_ref_ = 2.13 Å, in-plane hexagonal spacing) and gold at 1/*d*_ref_ = 0.426 Å^-1^ (*d*_ref_ = 2.35 Å, {111}). The peak position *r* (in Fourier pixels) was taken from the maximum of the resulting 1D radial intensity profile. For an N × N image region with pixel size p (Å/pixel), a peak at radial distance *r* corresponds to spatial frequency r/(N × p) Å^−1^. Equating this with 1/d gives:

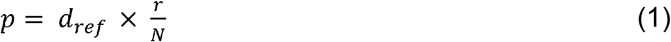

where *d*_ref_ is the known lattice spacing (Å), *d* is the measured radial peak position (Fourier pixels), and *N* is the image dimension (pixels).

### 2.3. Correction for elliptical magnification distortion

Anisotropic magnification or specimen tilt displaces Fourier-space lattice peaks from a circular arrangement to an elliptic arrangement; the resulting axis ratio carries direct information about either projector-lens anisotropy or stage tilt. WebCalEM exposes the ellipse-fitting algorithm of Yu *et al*. (Yu *et al*., 2016) as an interactive analysis option to recover the un-distorted pixel size. Quantitative validation of the recovered axis ratio against an independently characterized tilt or anisotropy is beyond the scope of this short communication; users with such datasets are encouraged to verify the recovery on their own systems.

### 2.4. Cross-standard consistency and measurement precision

No ground-truth microscope pixel size was available for direct accuracy assessment, so cross-standard consistency was used as a proxy. The agreement between pixel sizes independently calibrated from gold and graphene oxide under identical microscope settings, and the deviation of calibrated values from manufacturer-reported nominal values, were both reported. Cross-standard consistency between the gold and graphene-oxide measurements provides a check against sample-dependent systematic errors such as inaccurate reference lattice values, ring-fitting biases that vary with peak sharpness, or radial detector distortions that act differently on rings at different radii. The consistency was quantified using the Bland–Altman comparison framework (Bland & Altman, 1986; Giavarina, 2015), reporting the per-condition difference Δ = mean(GO) − mean(AU) with its 95% confidence interval, and summarizing across the ten (microscope, magnification) conditions as bias (mean Δ) and 95% limits of agreement (LoA) (mean Δ ± 1.96 × SD(Δ)). Given the modest number of condition-level pairs (n = 10), a small-sample-corrected multiplier was also computed to evaluate confidence bounds for the LoA (Carkeet, 2015). Precision was assessed by calibrating pixel sizes from 15–50 independent image regions at each magnification setting and computing the standard deviation of these measurements across different grid squares and imaging sessions for both calibration standards.

### 2.5. Web application implementation

WebCalEM is distributed in two interchangeable forms. The primary deliverable is a client-side written in JavaScript, delivered as a single self-contained HTML file. The app can be run directly in a browser from a local copy with no internet connection—a configuration well suited to isolated core-facility microscope workstations. A functionally equivalent server-side reference implementation, built in Python, is also distributed in the codebase and was used to benchmark the browser numerical core. Detailed UI implementation is discussed in section 3.3.(The Shiny development team)(Inc, 2015)

## 3. Results

### 3.1. Gold and graphene oxide yield consistent calibrations across magnifications and instruments

A key question for any internal-standard approach is whether different crystalline materials produce consistent results under identical imaging conditions. We addressed this by independently calibrating pixel sizes from gold and graphene oxide regions of the same grid (15–50 per magnification, depends on distribution of test specimens on collected micrographs) at five magnification settings on both Krios and Arctica microscopes (Fig. 1). Across all 10 conditions, mean pixel sizes from the two standards differed by less than 0.3% of the nominal value, with absolute measurement uncertainties on the order of 10^−3^ Å/pixel for both materials. The largest discrepancy between standards occurred on the Arctica at the 0.970 Å/pixel nominal setting (150,000×), where graphene oxide measurements ran ∼0.26% below gold; the two means still agreed within their combined measurement uncertainties at this condition.

**Figure 1.**
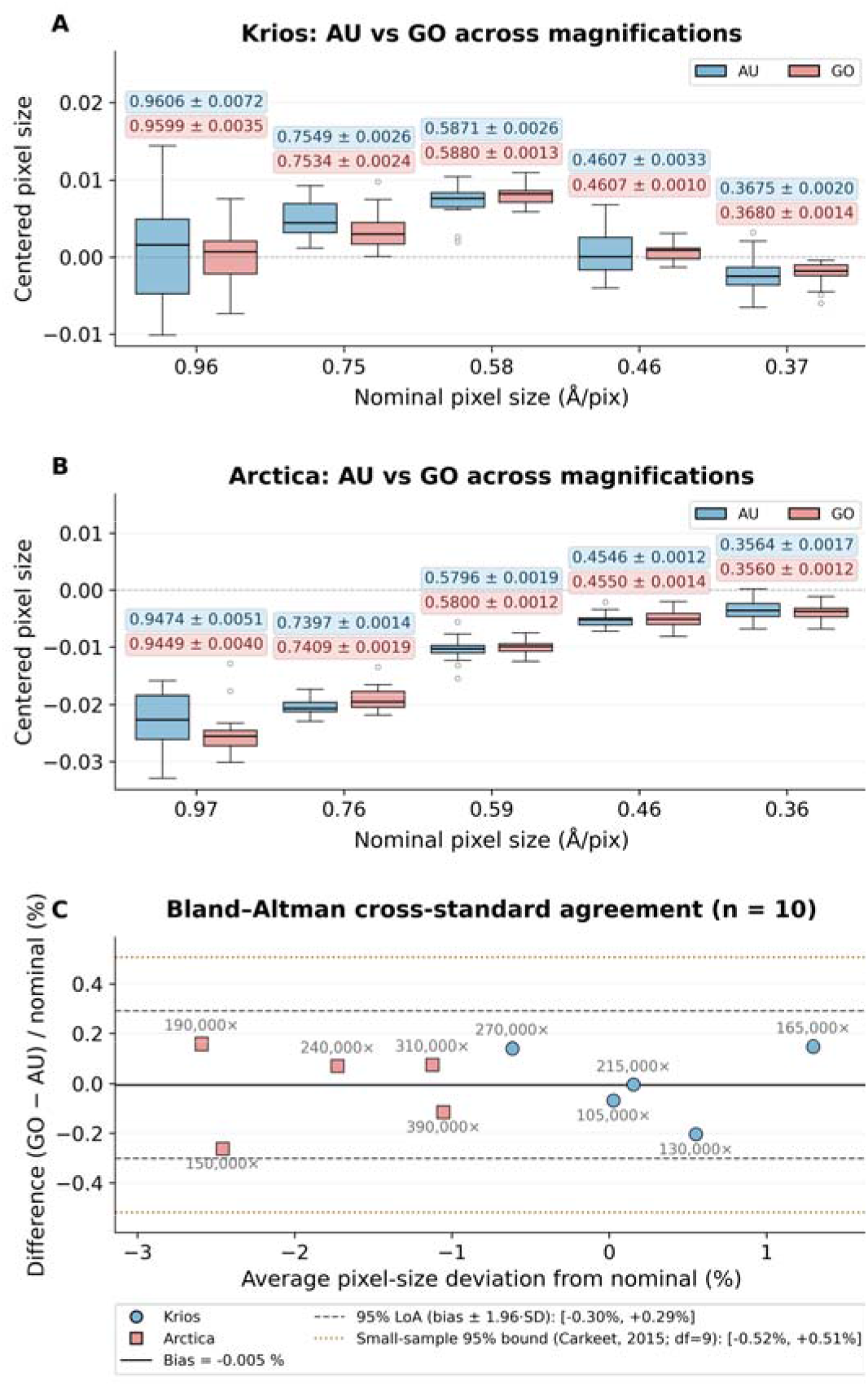
Centered pixel size distributions by nominal magnification and specimen type. Box plots show the distribution of calibrated pixel sizes (after subtracting the nominal value (dashed line)) for gold (blue) and graphene oxide (red) across five magnification settings on (A) Titan Krios G3 and (B) Talos Arctica G2 microscopes. (C) Bland–Altman comparison of GO and AU calibrations expressed as percent of nominal. Bias is −0.005% of nominal; 95% LoA are [−0.30%, +0.29%].

A Bland–Altman analysis across the ten conditions yielded an essentially zero overall bias of −0.005% of nominal with 95% LoA of [−0.30%, +0.29%]. Applying a t-based small-sample correction (Carkeet, 2015) for N=10 gave confidence bounds for the LoA of [−0.58 %, +0.56 %] of nominal, indicating sub-percent cross-standard discrepancy between gold and graphene oxide across the tested conditions. On both instruments, calibrated pixel sizes deviated systematically from nominal. The Arctica read uniformly low—about −2.6% at the lowest magnifications, shrinking to −1% at the highest. The Krios was non-monotonic: near nominal at the lowest magnification, peaking at +1.3% mid-range, and crossing to −0.6% at the highest. Both standards reproduced these instrument-specific deviations to within ∼0.3% at every condition, indicating they are not calibration-method artifacts and reinforcing the need for empirical, per-instrument calibration.

### 3.2. Retrospective calibration of archived datasets

Internal crystalline standards present on sample grids enable recalibration of previously collected data without recollection. We demonstrated this using archived micrographs from two amyloid-beta fibril datasets (unpublished)—ex vivo amyloid dataset collected at 105,000× on the Titan Krios G3, and in vitro amyloid data, collected at 190,000× on the Talos Arctica G2. For these datasets, GO–coated grids had been chosen to increase sample concentration rather than for calibration purposes. By selecting background regions of these micrographs, we were able to retrospectively calibrate the pixel size. The recovered values showed excellent agreement with those obtained from dedicated test micrographs at the corresponding magnification settings (Fig. 2). These examples demonstrate the feasibility of retrospective calibration when archived micrographs contain sufficiently visible crystalline support reflections in analyzable background regions.

**Figure 2.**
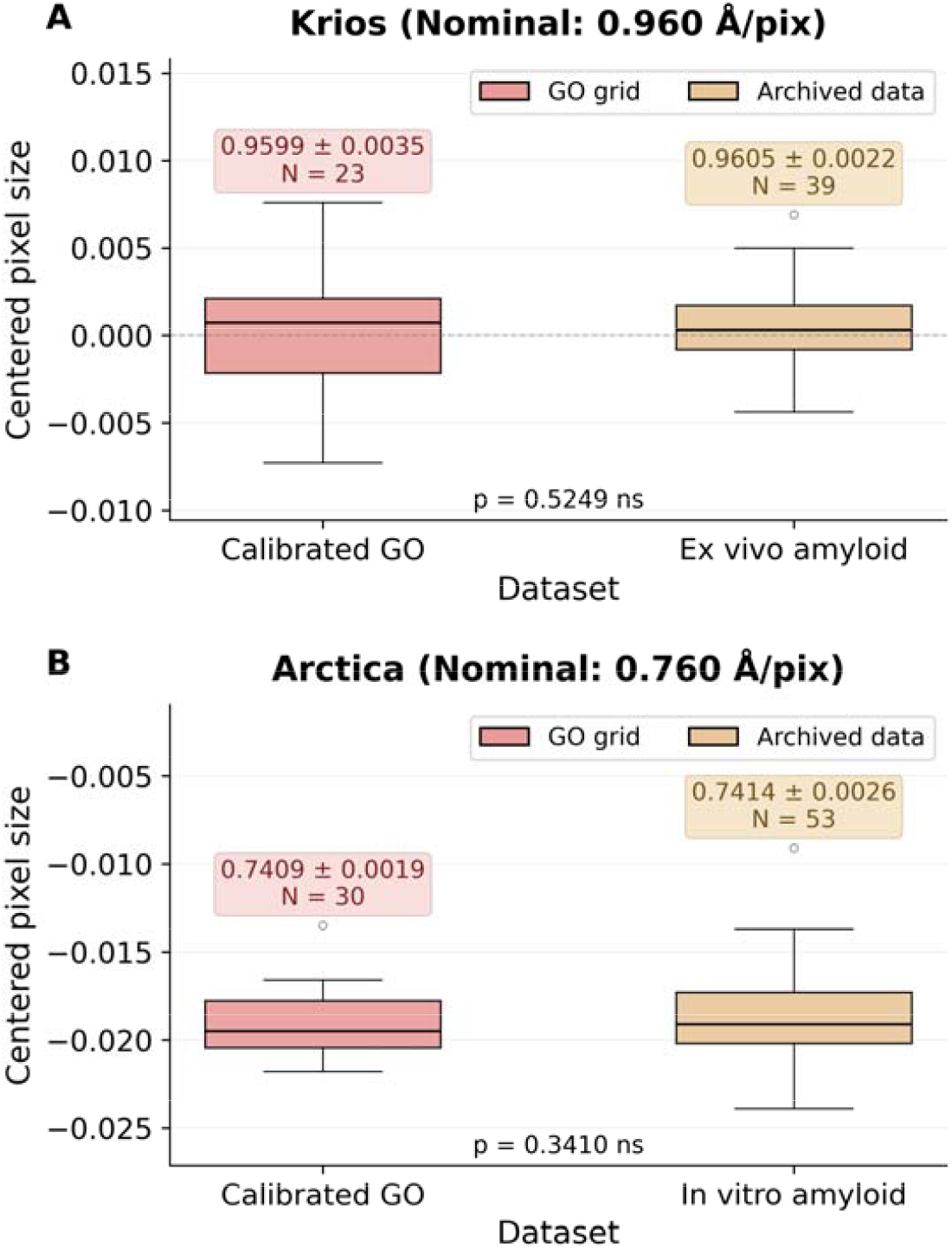
Retrospective per-region calibration on archived amyloid-β fibril datasets compared with calibrations from dedicated test micrographs at matching magnification settings. (A) Titan Krios G3 at 105,000× (nominal 0.96 Å/pix): new GO calibration versus the ex vivo amyloid dataset. (B) Talos Arctica G2 at 190,000× (nominal 0.76 Å/pix): new GO calibration versus the in vitro amyloid dataset. p-values are from independent-samples t-tests on the centered pixel-size distributions.

### 3.3. Web App Implementation

WebCalEM provides an integrated interface for pixel size calibration organized into three functional panels (Fig. 3). The image panel (blue) allows users to upload cryo-EM micrographs and interactively select regions of interest for analysis. The FFT analysis panel (yellow) offers both 1D radial profile and the 2D power spectrum. Peak detection is performed automatically in both views—in the 1D profile, the software identifies the radial maximum near the expected lattice frequency, while in the 2D spectrum, lattice peak positions are detected, and an ellipse is fitted automatically for distortion estimation. Users can manually adjust any of these results if needed. Radial and angular sampling frequencies are also adjustable, and a focused heatmap around the selected peak provides visual confirmation of the localization. The results panel (green) displays the calibrated pixel size for the current measurement and maintains a running table of all measurements within the session, automatically computing the group statistics for each nominal magnification. An interactive scatter plot visualizes the distribution of measurements centered by their nominal values, enabling users to assess calibration consistency immediately. Saved measurements can be reloaded and edited by clicking a row in the table, which restores the original image and region for refinement before being written back to the table. Results export contains a measurements CSV, a grouped-statistics CSV per nominal magnification, and the statistics scatter plot as PNG—suitable for direct integration into downstream processing pipelines. An auto-sample mode is also provided for batch use: the user selects a region count and region size, points the app at one or more images, and the app draws regions automatically and records the per-region pixel size for each. A detailed walk-through is released along with the codebase.

**Figure 3.**
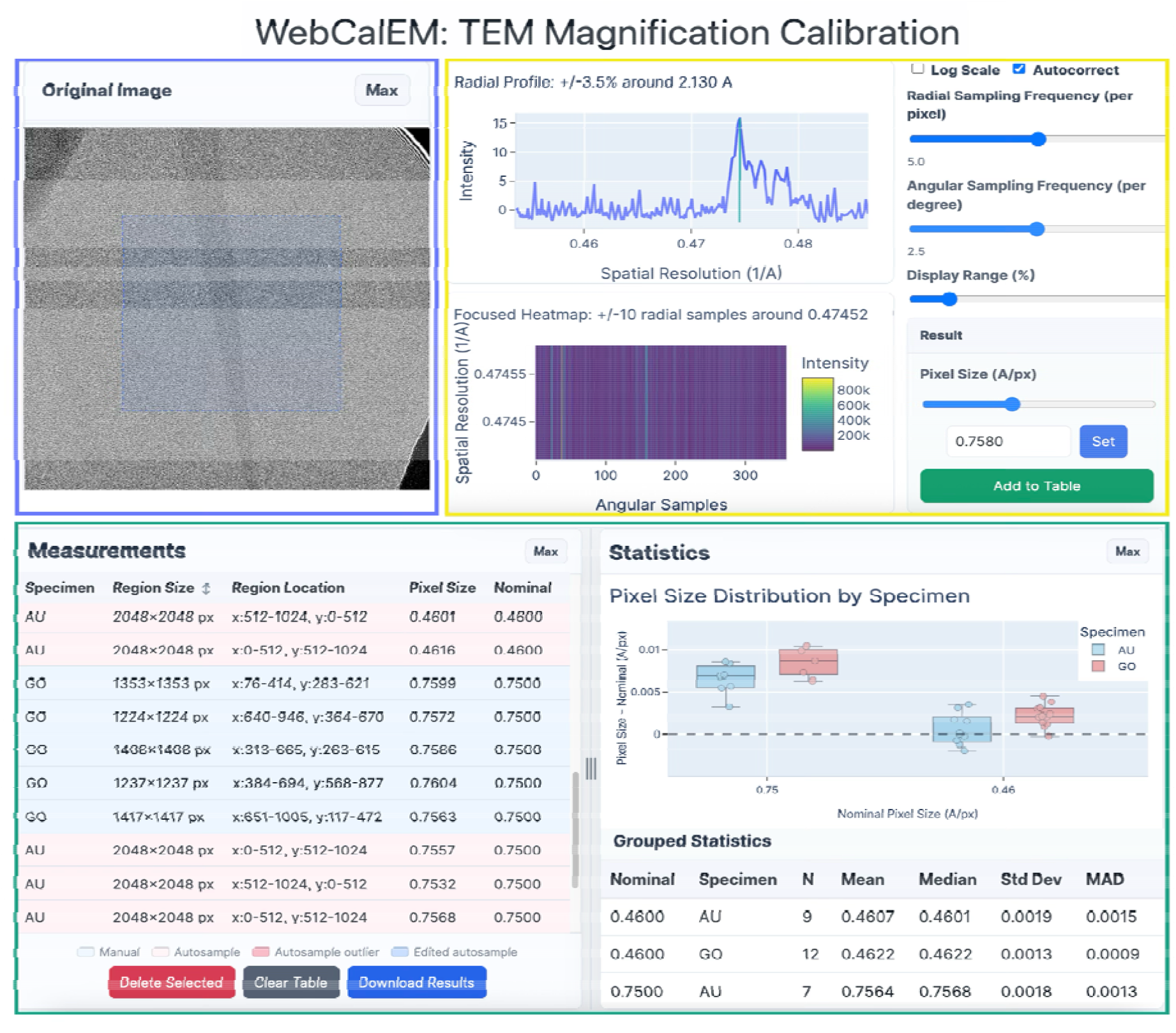
WebCalEM App user interface.

## 4. Discussion

Although both standards yielded statistically equivalent calibrations across the conditions we tested, the two materials differ in practical respects that affect routine use. The graphene-oxide hexagonal lattice produces a single, well-isolated 2.13 Å reflection that is unambiguous to pick in the 1D radial profile, whereas UltrAuFoil grids expose two closely spaced gold reflections at 2.35 Å and 2.48 Å—separated by only ∼5% in reciprocal space. This is wide enough to resolve the two rings most of the time, but the two bands are easily confused at lower nominal magnifications, where the rings sit closer together in Fourier pixels. GO peak picking was therefore more reliable than AU peak picking across the magnification range we tested, an observation consistent with the slightly tighter per-condition GO uncertainties we obtained (median ∼0.31% of nominal for GO versus ∼0.46% for AU) (Fig. 1). The two materials also differ in the nature of their elliptical distortion. When gold is used, the apparent ellipticity of the 2D power spectrum is dominated by global effects—projector-lens magnification anisotropy, stage tilt, and gradients in ice thickness—that are largely invariant across imaging regions on a single grid, so the ellipse fit can be evaluated once for the entire dataset and applied uniformly. Graphene-oxide flakes, however, are not always flat on the support film; physical out-of-plane tilt of the flake introduces a local ellipticity that depends on the specific region selected, so with GO the ellipse fit needs to be re-evaluated per region rather than per dataset. WebCalEM exposes interactive per-region ellipse fitting for exactly this reason.

In general, WebCalEM reaches the precision band of established command-line lattice-calibration tools (e.g., <0.5% of nominal for magCalEM (Dickerson *et al*., 2024)) while removing the installation and command-line barriers that have historically kept routine calibration out of standard cryo-EM workflows. The auto-sample mode introduced here already extends the workflow to batch calibration of pre-acquired datasets; the natural next step is to aggregate per-instrument calibrations into a shared community database—turning pixel-size calibration from an occasional procedure into a routine, field-wide quality-assurance practice.

## 5. Conclusion

We have developed WebCalEM, a browser-based tool that makes lattice-based pixel-size calibration a routine step in the cryo-EM workflow. Delivered as a single self-contained HTML file that runs entirely client-side—with no server, installation, or internet connection required after first load—it is deployable in any computational setting at a standard facility. The application supports multiple lattice standards, manual and auto-sampled per-region measurements, and automated statistical aggregation, while matching the precision of established command-line calibration tools. Widespread adoption should improve the accuracy and cross-comparability of cryo-EM structures across the field.

## Code and software availability

WebCalEM is free and open-source software developed by the Jiang Lab at Penn State University, with the source code on GitHub at https://github.com/jianglab/WebCalEM. The client-side single-file web app is hosted at https://jianglab.github.io/WebCalEM/. A detailed walkthrough is released as part of the codebase. No new structures, models, or maps were deposited as part of this work; PDB, EMDB, and EMPIAR deposition are therefore not applicable to this Methods Communication. Source micrographs supporting the cross-standard analysis are available from the corresponding author upon request.

## Author Contributions

L.D. developed the web application, collected cryo-EM data, performed all calibration and statistical analyses, and wrote the manuscript. Z.W., S.H.C., and S.L. contributed to cryo-EM data collection. G.C. prepared the GO-coated grids. N.F. collected the data used for retrospective calibration. W.J. conceived, supervised the project and revised the manuscript. All authors discussed the results and contributed to the final manuscript.

## Acknowledgements/Funding

This work was supported in part by NIH grants R01AG071177, RF1NS110437, R01AG091375, and Penn State startup fund (WJ). We would like to acknowledge the Huck Institutes’ Cryo-Electron Microscopy (Facility (RRID: SCR_024456) for use of the Thermo Fisher Titan Krios and Talos Arctica.

## References

Bland, J. M. & Altman, D. G. (1986). Lancet 1, 307–310.

Carkeet, A. (2015). OVS 92, 10.1097/OPX.0000000000000513.

Dickerson, J. L., Leahy, E., Peet, M. J., Naydenova, K. & Russo, C. J. (2024). Ultramicroscopy 256, 113883.

Giavarina, D. (2015). Biochem Med (Zagreb) 25, 141–151.

Han, Y., Fan, X., Wang, H., Zhao, F., Tully, C. G., Kong, J., Yao, N. & Yan, N. (2020). Proc. Natl. Acad. Sci. U.S.A. 117, 1009–1014.

Heimowitz, A., Andén, J. & Singer, A. (2020). Ultramicroscopy 212, 112950.

Inc, P. T. (2015). https://plot.ly.

Lövestam, S. & Scheres, S. H. W. (2022). Faraday Discuss. 240, 243–260.

MAG*I*CAL™ Calibration Standard for TEM Electron Microscopy Sciences.

McCaffrey, J. P. & Baribeau, J.□M. (1995). Microscopy Res & Technique 32, 449–454.

Meyer, J. C., Geim, A. K., Katsnelson, M. I., Novoselov, K. S., Booth, T. J. & Roth, S. (2007). Nature 446, 60–63.

Naydenova, K., Jia, P. & Russo, C. J. (2020). Science 370, 223–226.

Naydenova, K. & Russo, C. J. (2022). Ultramicroscopy 232, 113396.

Palovcak, E., Wang, F., Zheng, S. Q., Yu, Z., Li, S., Betegon, M., Bulkley, D., Agard, D. A. & Cheng, Y. (2018). Journal of Structural Biology 204, 80–84.

Penczek, P. A. (2010). Methods Enzymol 482, 73–100.

Penczek, P. A. (2020). IUCrJ 7, 995–1008.

Rohou, A. & Grigorieff, N. (2015). Journal of Structural Biology 192, 216–221.

Russo, C. J. & Passmore, L. A. (2014). Science 346, 1377–1380.

Sader, K., Stopps, M., Calder, L. J. & Rosenthal, P. B. (2013). J Struct Biol 183, 531–536.

Scheres, S. H. W. (2020). Acta Crystallogr D Struct Biol 76, 94–101.

The Shiny development team Shiny for Python.

Tiwari, S. P., Chhabra, S., Tama, F. & Miyashita, O. (2020). J. Chem. Inf. Model. 60, 2570–2580.

Wang, J., Liu, J., Gisriel, C. J., Wu, S., Maschietto, F., Flesher, D. A., Lolis, E., Lisi, G. P., Brudvig, G. W., Xiong, Y. & Batista, V. S. (2022). Journal of Structural Biology 214, 107902.

Wasilewski, S., Karelina, D., Berriman, J. A. & Rosenthal, P. B. (2012). Journal of Structural Biology 180, 243–248.

Yu, G., Li, K., Liu, Y., Chen, Z., Wang, Z., Yan, R., Klose, T., Tang, L. & Jiang, W. (2016). Journal of Structural Biology 195, 207–215.

